# What the structural-functional connectome reveals about brain aging: The key role of the fronto-striatal-thalamic circuit and the rejuvenating impact of physical activity

**DOI:** 10.1101/183939

**Authors:** P. Bonifazi, A. Erramuzpe, I. Diez, I. Gabilondo, M.P. Boisgontier, L. Pauwels, S. Stramaglia, S.P. Swinnen, J.M. Cortes

## Abstract

Physiological ageing affects brain structure and function impacting its morphology, connectivity and performance. However, at which extent brain-connectivity metrics reflect the age of an individual and whether treatments or lifestyle factors such as physical activity influence the age-connectivity match is still unclear. Here, we assessed the level of physical activity and collected brain images from healthy participants (N=155) ranging from 10 to 80 years to build functional (resting-state) and structural (tractography) connectivity matrices that were combined as connectivity descriptors. Connectivity descriptors were used to compute a maximum likelihood age estimator that was optimized by minimizing the mean absolute error. The connectivity-based estimated age, i.e. the brain-connectome age (BCA), was compared to the chronological age (ChA). Our results were threefold. First, we showed that ageing widely affects the structural-functional connectivity of multiple structures, such as the anterior part of the default mode network, basal ganglia, thalamus, insula, cingulum, hippocampus, parahippocampus, occipital cortex, fusiform, precuneus and temporal pole. Second, our analysis showed that the structure-function connectivity between basal ganglia and thalamus to orbitofrontal and frontal areas make a major contribution to age estimation. Third, we found that high levels of physical activity reduce BCA as compared to ChA, and vice versa, low levels increment it. In conclusion, the BCA model results highlight the impact of physical activity and the key role played by the connectivity between basal ganglia and thalamus to frontal areas on the process of healthy aging. Notably, the same methodology can be generally applied both to evaluate the impact of other factors and therapies on brain ageing, and to identify the structural-functional brain connectivity correlate of other biomarkers than ChA.

## Introduction

Ageing may be defined as a time-dependent decline involving an accumulation of changes at the biological, psychological, and social level^1^. Interestingly, individuals with the same chronological age (ChA) exhibit different trajectories of age-related biological deterioration, as measured by biomarkers of functional performance, tissue integrity and metabolic health^2,3^. This mismatch reflects two different concepts for ageing. One is ChA, calculated as the time running since birth, whereas the other is the biological age, which, irrespective of birth year, is based on the level of biological functioning at a given time. The mismatch between chronological and biological ageing has gained major scientific interest in the past years due to the potential implication on health and disease of age-related molecular, genetic, cellular and organ-specific dynamics and their genetic, epigenetic, and environmental modulators^4^. In fact, it is well established that ageing is one of the main risk factors for most late-onset diseases such as cancer, cardiovascular disease, diabetes and neurodegenerative diseases^5^.

In terms of biological brain ageing, psychophysical, neuropsychological and physiological studies support the fact that brain functional performance declines with age, with specific impact on cognition (long-term and working memory, executive functions, conceptual reasoning and processing speed)^6,7^, mood (anxiety and depression)^8^, circadian behaviours (disruption of amplitude and period length) and sleep cycle (poor sleep quality and delayed sleep onset latency)^9^.

These changes in brain performance occur in parallel with well-established age-related macrostructural and microstructural brain variations. At microstructural level, age has been associated with alterations in synaptic structures (decreased synaptic density and synaptic terminals), aggregation of abnormal proteins outside and inside neurons such as plaques and tangles, reduced neurogenesis and synaptic plasticity, abnormal increase of astrocytes and oligodendrocytes, altered myelination and reduction of nerve growth factor concentration^10–12^.

The effects of age on the reduction of the number of neurons in the brain has been under debate, with several post-mortem human and primate studies supporting the fact that cortical cell count remains unchanged^13^ and that neuronal shrinking (rather than cell loss) is the main process underlying brain atrophy. However, at the macroscopic level, both global and regional atrophies are the best-reported characteristics of the ageing brain, as supported by several post-mortem and MRI studies. Neuroimaging studies have shown that the overall brain volume varies with age in an “inverted-U” fashion consisting of an increase of about 25% from childhood to adolescence, then remaining constant for about three decades to finally decay down to childhood size in late ages^14^. This pattern of age-related brain atrophy in the elderly has been associated with the deterioration of cognitive performance in the healthy population^15^. Of note, age-related grey and white matter brain atrophies are not homogeneous, with higher atrophy rates observed in white matter as compared to cortical grey matter^16,17^ and regionally, with more prominent atrophy in prefrontal and parietal cortices^18–20^ and hippocampus^21^. In contrast, the volume of the cerebrospinal space (ventricles, fissures, and sulci) increases with age^16^.

In line with macrostructural MRI correlates of brain ageing, modern techniques such as diffusion tensor imaging (DTI) have facilitated the in-vivo inspection of age-related microstructural changes of the brain^22,23^, supporting histological findings and revealing regional patterns and dynamics of structural connectivity (SC) degeneration, a phenomenon postulated to lead to cortical disconnection with loss of functional integration of neurocognitive networks^7,24^. Several DTI studies in normal ageing support that white matter atrophy is associated with a widespread microstructural degeneration of white matter fibers, with changes predominantly affecting frontal tracts^24^, and gradually extending to posterior tracts^25^, a pattern that inverts the sequence of myelination during early development and supports the “last-in-first-out principle” for white matter deterioration along the lifespan.

The development of resting and task-based functional MRI (fMRI) has provided in-vivo functional insights into the observed age-related atrophy and SC brain disconnection, consistently showing age-related regional changes in the patterns of brain activation, with decreased activity in the occipital lobe and increased activity in the frontal lobe across a variety of tasks^7^. Functional connectivity (FC) studies with resting state fMRI have gone a step further demonstrating that ageing not only induces regional brain activity changes but also a decrease in functional connectivity of large-scale brain networks, specifically between anterior and posterior hubs, including superior and middle frontal gyrus, posterior cingulate, middle temporal gyrus, and the superior parietal region^26,27^.

The combination of SC and FC analyses by complex network approaches have led to the conceptualization of brain networks as a connectome^28,29^, and its correlates with age and diseases has gained major attention in fundamental neuroscience^30^. Complex network approaches have highlighted the key role played by several network descriptors in ageing and brain diseases, such as network hubness, hub integrity, network modularity and hierarchical organization of networks. In terms of connectome modularity, it has been shown that functional network modularity (segregation) decreases with age^31,32^, a mechanism supporting the loss of functional specialization of certain key brain networks known to be involved in the cognitive domains affected by ageing along the lifespan^33^.

Moreover, combined SC and FC analyses have suggested that not only segregation (i.e., network modularity) decreases with age but integration (i.e., node efficiency) increases^34^ in a counterbalanced manner assuring network efficiency along lifespan. Others, however, have suggested that small-worldness and network modularity remain stable along the lifespan, despite a considerable reduction in streamline number ^35^. Analyses of longitudinal data showed that age variations affecting FC did not correspond with the variations in SC^36^, highlighting that FC and SC were affected by age in a more independent manner than previously thought. Conversely, the age variations in FC and SC between areas participating in the DMN were highly correlated with each other.

The combined FC and SC analysis also revealed the key role of structural deterioration of the cortical-subcortical connections in the integration of several resting state networks and performance on cognitive tasks, such as those involving executive functions, processing speed and memory^37^. By calculating node-degree distributions, other studies have shown a reduction with age in the connectivity degree of network hubs ^38^, supporting the theory that the alteration of network hubs underlies brain physiological ageing as well as a plethora of different brain pathologies^30^.

Recently, new computational strategies for analysing the dynamics of brain atrophy (such as machine learning) have introduced the concept of brain-predicted ageing, which facilitates the quantification of the mismatch between age-related brain atrophy and ChA. Deviations of brain-predicted ageing from healthy brain ageing have been described for several brain diseases, including traumatic brain injury^39^, mild cognitive impairment and Alzheimer’s disease^40^, HIV infection^41^ and schizophrenia^42^. A critical issue in the study of brain biological ageing by computational neuroimaging is the selection of an adequate approach with the highest robustness and precision for the quantification of the mismatch between chronological and brain biological ages. The combined rather than separate analysis of SC and FC has shown to provide a better estimation of ChA^43^.

In line with this effort to improve ChA estimation approaches of SC and FC connectivity analyses, it is also of major importance to consider the role of potential epigenetic and environmental modulators of brain biological ageing on MRI derived measures. We focus here on physical activity (PA), that has been shown to reduce disability^44^, morbidity^45,46^ and mortality^47–49^. Regarding the effects of PA on brain biological functioning, it has been demonstrated in animal models that PA has a beneficial effect on learning and memory function by inducing neuroprotection, decreasing oxidative stress and increasing cerebral blood perfusion, which in turn influence angiogenesis, synaptogenesis and neurogenesis to eventually lead to memory improvement detected at the behavioral level^50^.

Although previous publications have addressed the study of brain ageing and the estimation of ChA by means of separate or combined analyses of SC and FC, none of the latter studies has proposed an optimal method that, applying complex network analysis of the structural-functional connectome, simultaneously identifies age-related brain changes and estimates brain age, considering the effect of potential key modulators of brain ageing such as PA.

In the present study, we have to key hypotheses; first, that the estimation of age is improved by combining FC and SC descriptors, and second, that higher and lower levels of PA might be associated with younger and older biological age, respectively. To test these hypotheses, following previous work^39–41,51–59^, we built an ageing data-driven model to estimate the ChA of participants based on SC and FC biomarkers and investigated the extent to which the level of PA mediates the participant’s brain biological age. Lastly, we discuss the general implications and applications of the described methodology.

## Material and Methods

### Participants

Participants were recruited in the vicinity of Leuven and Hasselt (Belgium) from the general population by advertisements on websites, announcements at meetings and provision of flyers at visits of organizations and public gatherings (PI: Stephan Swinnen). A sample of N=155 healthy volunteers (81 females) ranging in age from 10 to 80 years (mean age 44.4 years, SD 22.1 years) participated in the study. All participants were righthanded, as verified by the Edinburgh Handedness Inventory. None of the participants had a history of ophthalmological, neurological, psychiatric or cardiovascular diseases potentially influencing imaging or clinical measures. Informed consent was obtained before testing. The study was approved by the local ethics committee for biomedical research, and was performed in accordance with the Declaration of Helsinki.

### Physical activity score

Physical activity (PA) was assessed using the International Physical Activity Questionnaire^60^ (IPAQ), which assesses PA undertaken across leisure time, domestic and gardening activities, and work-related and transport-related activities. The specific types of activity were classified into three categories: walking, moderate-intensity activities, and vigorous-intensity activities. Frequency (days per week) and duration (time per day) were collected separately for each specific activity category. The total score used to describe PA was computed as the weighted summation of the duration (in minutes) and frequency (days) for walking, moderate-intensity, and vigorous-intensity activity. Each type of activity was weighted by its energy requirements defined in Metabolic Equivalent of Task (MET): 3.3 METs for walking, 4.0 METs for moderate physical activity, and 8.0 METs for vigorous physical activity^61^.

### Imaging acquisition

Magnetic resonance imaging (MRI) scanning was performed on a Siemens 3T Magnetom Trio MRI scanner with a 12-channel matrix head coil.

#### Anatomical data

A high resolution T1 image was acquired with a 3D magnetization prepared rapid acquisition gradient echo (MPRAGE): repetition time [TR] = 2300 ms, echo time [TE] = 2.98 ms, voxel size = 1 × 1 × 1.1mm^3^, slice thickness = 1.1 mm, field of view [FOV] = 256 × 240mm^2^, 160 contiguous sagittal slices covering the entire brain and brainstem.

#### Diffusion Tensor Imaging

*A DTI* SE-EPI (diffusion weighted single shot spin-echo echo-planar imaging) sequence was acquired with the following parameters: [TR] = 8000 ms, [TE] = 91 ms, voxel size = 2.2 × 2.2 × 2.2 mm^3^, slice thickness = 2.2 mm, [FOV] = 212 × 212 mm^2^, 60 contiguous sagittal slices covering the entire brain and brainstem. A diffusion gradient was applied along 64 non-collinear directions with a b value of 1000 s/mm^2^. Additionally, one set of images was acquired without diffusion weighting (*b= 0* s/mm^2^).

*Resting state functional data* was acquired over a 10 minute session using the following parameters: 200 whole-brain gradient echo echo-planar images with [TR/TE] = 3000/30 ms, [FOV] = 230 × 230mm^2^, voxel size = 2.5 × 2.5 × 3.1mm^3^, 80 × 80 matrix, slice thickness = 2.8 mm, 50 sagittal slices, interleaved in descending order.

### Imaging preprocessing

#### Diffusion Tensor Imaging

We applied DTI preprocessing similar to previous work^62–66^ using FSL (FMRIB Software Library v5.0) and the Diffusion Toolkit. First, an eddy current correction was applied to overcome the artefacts produced by variation in the direction of the gradient fields of the MR scanner, together with the artefacts produced by head motion. To ensure that correlations with age were not due to differences in head motion (ie., to correct for the effect that older people move more), the average motion of each participant was used as a covariate of non-interest in the statistical analyses. In particular, the participant’s head motion was extracted from the transformation applied at the eddy current correction step, from every volume to the reference volume (the first volume, *b=0*). The motion information was also used to correct the gradient directions prior to the tensor estimation. Next, using the corrected data, a local fitting of the diffusion tensor was applied to compute the diffusion tensor model for each voxel. Next, a Fiber Assignment by Continuous Tracking (FACT) algorithm was applied^67^. We then computed the transformation from the Montreal Neurological Institute (MNI) space to the individual-participant diffusion space and projected a high resolution functional partition to the latter, composed of 2514 regions and generated after applying spatially constrained clustering to the functional data^68^. This allowed building 2514 x 2514 SC matrices, each per participant, by counting the number of white matter streamlines connecting all region pairs within the entire 2514 regions dataset. Thus, the element matrix (i,j) of SC is given by the streamlines number between regions i and j. SC is a symmetric matrix, where connectivity from i to j is equal to that from j to i. Exclusion criteria was based on not having the average head motion higher than the mean + 2 standard deviation. None of the participants were excluded based on this constraint.

#### Functional MRI

We applied resting fMRI preprocessing similar to previous work^62–64,66,69,70^ using FSL and AFNI (http://afni.nimh.nih.gov/afni/). First, slice-time correction was applied to the fMRI dataset. Then each volume was aligned to the middle volume to correct for head motion artefacts. Next, all voxels were spatially smoothed with a 6 mm full width at half maximum (FWHM) isotropic Gaussian kernel and after intensity normalization, a band pass filter was applied between 0.01 and 0.08 Hz^71^ followed by the removal of linear and quadratic trends. We next regressed out the motion time courses, the average cerebrospinal fluid (CSF) signal, the average white-matter signal and the average global signal. Finally, the functional data was spatially normalized to the MNI152 brain template, with a voxel size of 3*3*3 mm^3^. In addition to head motion correction, we performed scrubbing, by which time points with framewise displacement higher than 0.5 were interpolated by a cubic spline^72^. Further, to correlate with age, we also removed the effect of head motion by using the global frame displacement as a non-interest covariate, as old participants moved more than the young, and this fact introduced *trivial* correlations with age. Finally, FC matrices were calculated by obtaining the pairwise Pearson correlation coefficient between the resting fMRI time series. Exclusion criteria was based on not having more than 20% of the time points with a frame wise displacement greater than 0.5. Two participants were finally excluded.

### Brain Hierarchical Atlas (BHA) and its robustness along lifespan

The aforementioned 2514 brain regions were grouped into modules using the Brain Hierarchical Atlas (BHA), recently developed^64^ and applied by the authors in a traumatic brain injury study^66^. The BHA is available to download at http://www.nitrc.org/projects/biocr_hcatlas/. A new Python version was developed during Brainhack Global 2017 - Bilbao can be downloading at **github, to be amended before submission**

The use of the BHA guarantees two conditions simultaneously: 1) That the dynamics of voxels belonging to the same module is very similar, and 2) that the voxels belonging to the same module are structurally wired by white matter streamlines; see in figure 1 the high correspondence between SC and FC modules. The BHA provides a multi-scale brain partition, where the highest dendrogram level *M=1* corresponded to all 2514 regions belonging to a single module, coincident with the entire brain, whereas the lowest level *M=2514* corresponded to 2514 separated modules, all of them composed of only one region.

**Figure 1:**
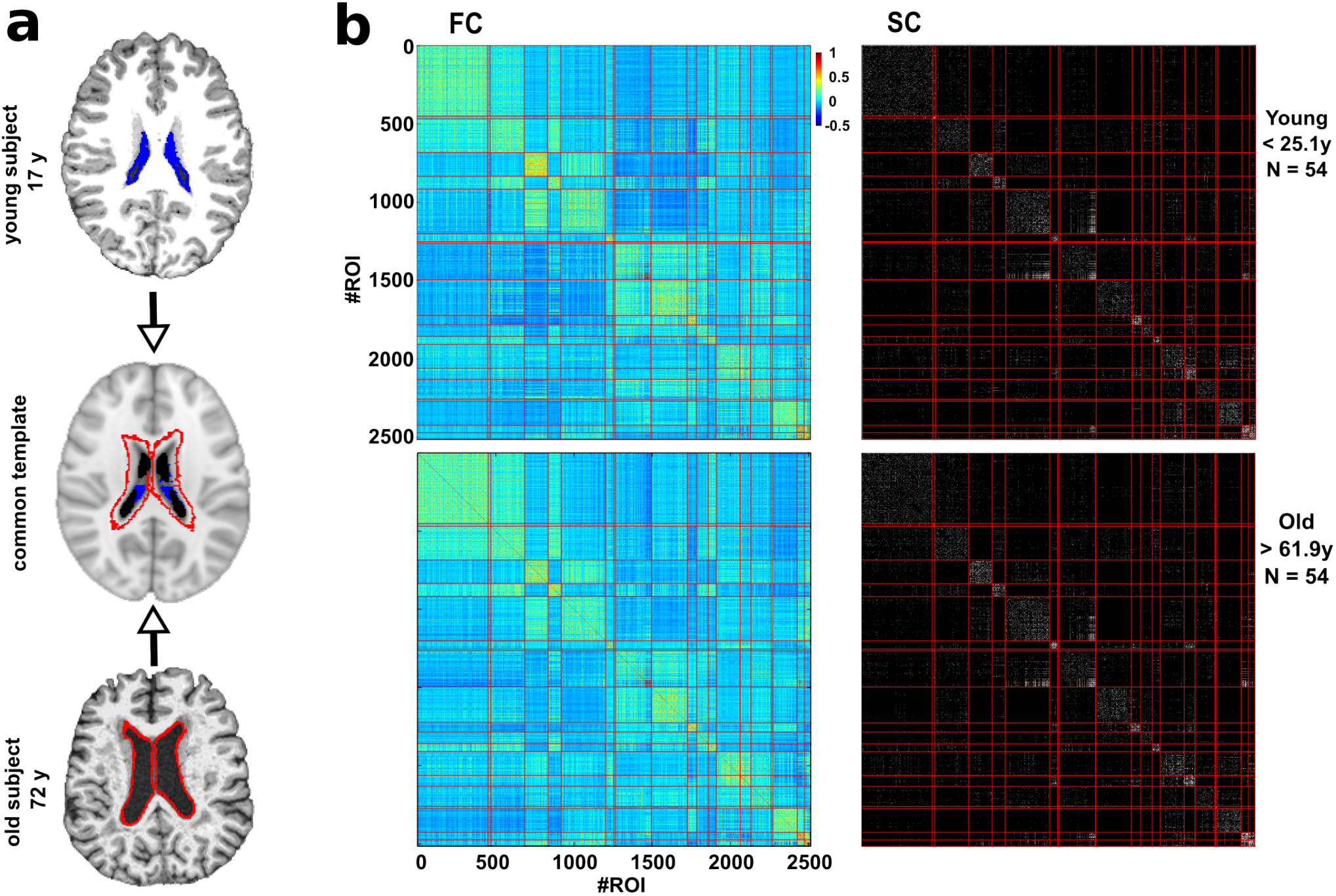
Robustness of the Brain Hierarchical Atlas along lifespan. **a:** Common template normalization (middle) for young (top) and old brains (bottom). Ventricle 3D segmentation has been performed for a young (17 y, filled in blue) and old participant (72y, contours marked in red). Both segmentations are superimposed onto the common population template (middle row). For the connectivity analysis, regions located within the volume defined by the biggest ventricle size across all the participants have been ignored to correct for trivial age-effects in the results of age estimation (i.e., to correct for the effect that older people have bigger ventricle volume). **b:** Brain hierarchical atlas (BHA) parcellation for young (top) and old (bottom) populations shows the strong correspondence between functional modules (depicted as yellow squares in the matrix diagonal of the functional connectivity matrix, FC) and structural modules (plotted in the SC matrix). FC and SC matrices are the result of averaging FC and SC individual matrices in two different populations, young (age < 25.1 y, *N=54* participants) and old (age > 61.9 y, *N=54* participants). Both connectivity FC and SC matrices have been reordered according to the BHA (here represented at the level of *M=20* modules). FC is defined by the pairwise Pearson correlation between rs-fMRI time series whilst SC is defined by the streamline counting between region pairs (here binarized just for illustration purposes).

It was also shown *in*^64^ that the hierarchical brain partition with M = 20 modules was optimal based on the cross-modularity index X. This index was defined as the geometric mean between the modularity of the structural partition (Q_s_), the modularity of the functional partition (Q_F_), and the mean Sorensen similarity between modules existing in the two structural and functional partitions (L_SF_).

### Labelling of anatomical regions

The anatomical representation of the 2514 brain regions were identified by using the Automated Anatomical Labelling^73^ (AAL) brain atlas. Therefore, the anatomical identification of the brain regions used in this work follow the labels appearing in the AAL atlas.

### Removal of regions affected by the increment of ventricular space along lifespan

Ventricular space increases along the lifespan in a manner that, after transforming all images to a common space, some regions surrounding the ventricular space for the younger population are occupied by the ventricular space of older participants. In order to remove this effect, we deleted these regions by (after projecting all images to the common space) searching for the participant with the highest ventricular volume, segmenting this space and treating it as mask to discard (for the connectivity analysis) all the regions within this space in all the participants. Figure 1a illustrates this procedure.

### Structure-function correlo-dendrograms of brain ageing

From both SC and FC matrices, we built the correlo-dendrogram (CDG) of brain ageing by correlating chronological age with the values of internal (intra-module) and external (inter-module) connectivity for each dendrogram level M of the BHA. In particular, four different classes of module descriptors were built per participant: functional internal connectivity (FIC), functional external connectivity (FEC), structural internal connectivity (SIC), and structural external connectivity (SEC) (figure 2). Given a brain module composed by a set of R regions, its associated FIC (SIC) was calculated as the sum of the functional (structural) weights of all the links between the elements of R, whilst FEC (SEC) was defined as the sum of the functional (structural) weights of all the links connecting the elements of R to other regions in the brain.

**Figure 2:**
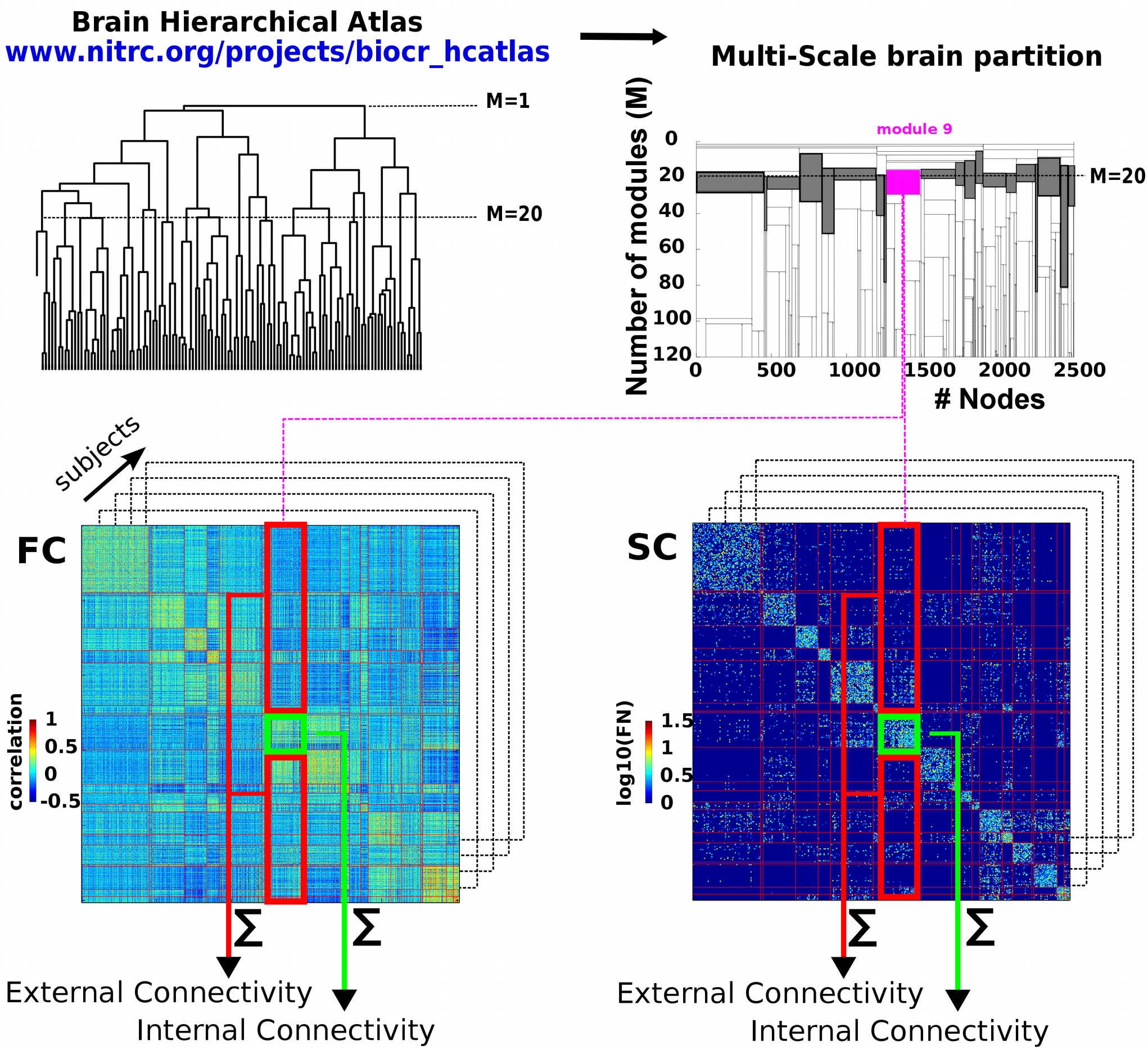
Schematic representation of brain connectivity descriptors. **Left-top:** First, we made use of BHA to define different modules resulting from a hierarchical agglomerative clustering. **Right-top:** The multiscale brain partition shows how modules divide when going down along the tree (here, we only considered the part of the tree that goes from 20 to 120 modules). The gray-colored modules represents the *M=20* brain partition. **Bottom:** For the tree level of *M=20* and for each participant, we calculated the structural/functional internal connectivity (green rectangle) and structural/functional external connectivity (red rectangle), by summing respectively the edge weights within and leaving out that module. The same procedure was applied for all the modules in all the 20<=M<=1000 levels of the tree.

One of the peculiarities of the BHA is that at each M level only one of the branches of the hierarchical tree divides in two, so at each level only 2 modules are new with respect to the (M-1) level (figure 2). Considering this characteristic and the fact that we started our analysis at the level of *M=20* and arrived up to *M=1000*, we established the Bonferroni significance threshold equal to 0.05/[20+2 ∗(1000 −20)] for the correlation between age and connectome measures (FIC, SIC, FEC, SEC).

To localize age-affected brain areas at both functional and structural levels (rather than separate FC or SC analyses), and thus obtaining a major benefit from the combination of functional and structural data, we searched for brain regions such that the p-value was smaller than the square root of the product of the individual structural p-value multiplied by the individual functional one. The value of the structure-function age correlation was calculated as the geometric mean of the two correlation values, one achieved by the functional descriptor and the other by the structural one.

A Python pipeline implementing this strategy can be downloaded at **github, to be amended before submission**

### Maximum likelihood estimator (MLE)

Let us define for each participant *n* the vector 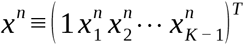 of *n* components, each one corresponding to a different connectivity descriptor (in principle any value of inter/intra module connectivity at any M level of the BHA calculated from either FC or SC), where *T* denotes the transpose operator. The estimated age for participant *n* was calculated by a linear combination of the descriptors, ie.,

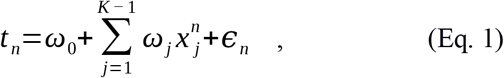

where ^*ε*^*n* is a zero mean Gaussian random variable with variance *σ*^2^ and *ω≡*(*ω*_0_ *ω*_1_*ω*_2_⋯ *ω*_*K-*1_)^*T*^ is the weight vector. For *P* different participants, using eq. (1), one can define the error function as

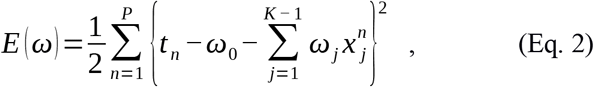

which allows to calculate the weight vector that minimizes the error function, that is, which is solution of *∇*_*ω*_ *E* (*ω*)=0 (first derivative with respect to *ω* equal to zero). Such a minimum defines precisely the Maximum Likelihood Estimator (MLE), which can be analytically solved^74,75^ and is given by:

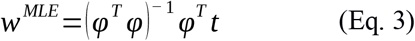

where -1 denotes the inverse of the matrix, *t≡*(*t*_1_*t*_2_*t*_3_⋯*t*_*P*_)^*T*^ is the vector of P age-participant estimations and *φ* is the so-called *design matrix*, ie.,

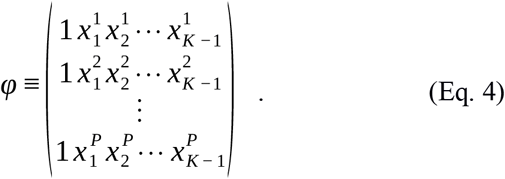

### Mean absolute error (MAE) and brain connectome age (BCA)

When the entire dataset is used to calculate *w*^*MLE*^, increasing the number of descriptors the error estimation function decreases *(the more descriptors, the better the estimation)*, but this strategy also provides a very high variance estimate, meaning that, when estimating the age using the *w*^*MLE*^ solution in a different dataset can produce a very high error. Splitting the entire dataset in training and testing sets solves this problem, well known as overfitting^76^.

To calculate the MAE, for each experiment we performed data splitting, by randomly choosing 75% of the dataset (N1=115) for training (i.e., for calculating the *w*^*MLE*^ solution) and the remaining 25% (N2=38) for testing (i.e., to calculate the MAE). As a metric for estimation quality, the MAE in the testing dataset was calculated as

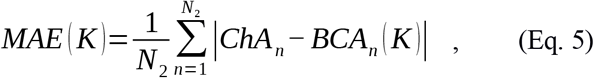

where || denotes absolute error and where we have defined the brain connectome age (BCA) for participant n as

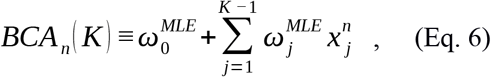

where *w*^*MLE*^ has been defined in Eqs. (3) and (4).

Remark that although in principle there were many potential descriptors (four classes –FIC, FEC, SIC and SEC– per module and number of modules M varying from 20 to 1000), finally only K of them were introduced into the MLE to estimate age. Therefore, and by construction, the MLE solution depends on K (see next subsection for the choice for the K descriptors).

### Optimization of the maximum likelihood estimation (MLE)

In order to get *the best model*, ie. the K descriptors that better estimate age, we optimized the MLE in the following way:

1. For K=1, we considered the descriptor that best correlated with ChA.
2. The K=2 descriptor was chosen among all the remaining ones by finding the descriptor such that after U=100 experiments of randomly choosing 75% of the dataset for training and 25% for testing, the mean MAE achieved by the two descriptors (the one found in stage 1 plus the new one) was minimal.
3. The K=3 descriptor was chosen among all the remaining ones by finding the descriptor such that after U=100 experiments of randomly choosing 75% of the dataset for training and 25% for testing, the mean MAE calculated with three descriptors (the previous two descriptors found in stage 2 plus the new one) was minimal.
4. Following this strategy, the curve MAE(K) has a minimum value as K increases, that defined the best model which has K descriptors.

This age estimation strategy has been implemented using *scikit-learn*, http://scikit-learn.org/stable/. The entire code can be downloaded at **github, to be amended before submission**

## Results

A population of N=155 healthy participants (81 female, 74 male) with age varying from 10 to 80 years (mean=44, standard deviation=22) was used for the study. Together with physical activity scores, triple acquisitions including anatomical, diffusion tensor and resting functional imaging were acquired for each participant.

We used the BHA Brain Hierarchical Atlas (BHA, see^64^ and Methods) with 2514 regions and calculated for each participant the weighted SC and FC connectivity matrices, representing respectively the region-pairwise streamline number and the region-pairwise Pearson correlation of resting-state activity time series.

We first verified the robustness of the BHA across lifespan. Figure 1 shows for two populations, one young (age < 25.1 y, N=54 participants) and other old (age > 61.9 y, N=54 participants), that the correspondence between SC modules and FC modules was preserved independently of participant age. This was quantified by assessing cross-modularity (X), obtaining X = 0.312 for the young population and X = 0.309 for the old, and therefore showing that cross-modularity was 99% preserved along the lifespan.

Next, we calculated for all levels M of the BHA (with 20<=M<=1000), four different module descriptors (figure 2): 1) the functional internal connectivity (FIC), 2) the functional external connectivity (FEC), 3) the structural internal connectivity (SIC) and 4) the structural external connectivity (SEC), that we used for building the correlo-dendrograms (CDG) that allowed to find the highest correlation between module connectivity and ChA, whilst maximizing the spatial resolution.

Figure 3 shows the structure-function CDGs, calculated as the geometric mean between the correlation value obtained from the functional descriptor and the correlation value obtained from the structural one. This allowed building one CDG based on external structure-function connectivity (figure 3a, associated to connections with longer fiber length) and a different one based on internal structure-function connectivity (figure 3b, associated to connections with shorter fiber length). Brain maps from the external CDG (figure 3a) and the internal CDG (figure 3b) were built assigning to each brain region the maximum absolute value of structural-functional correlation across all BHA levels.

**Figure 3.**
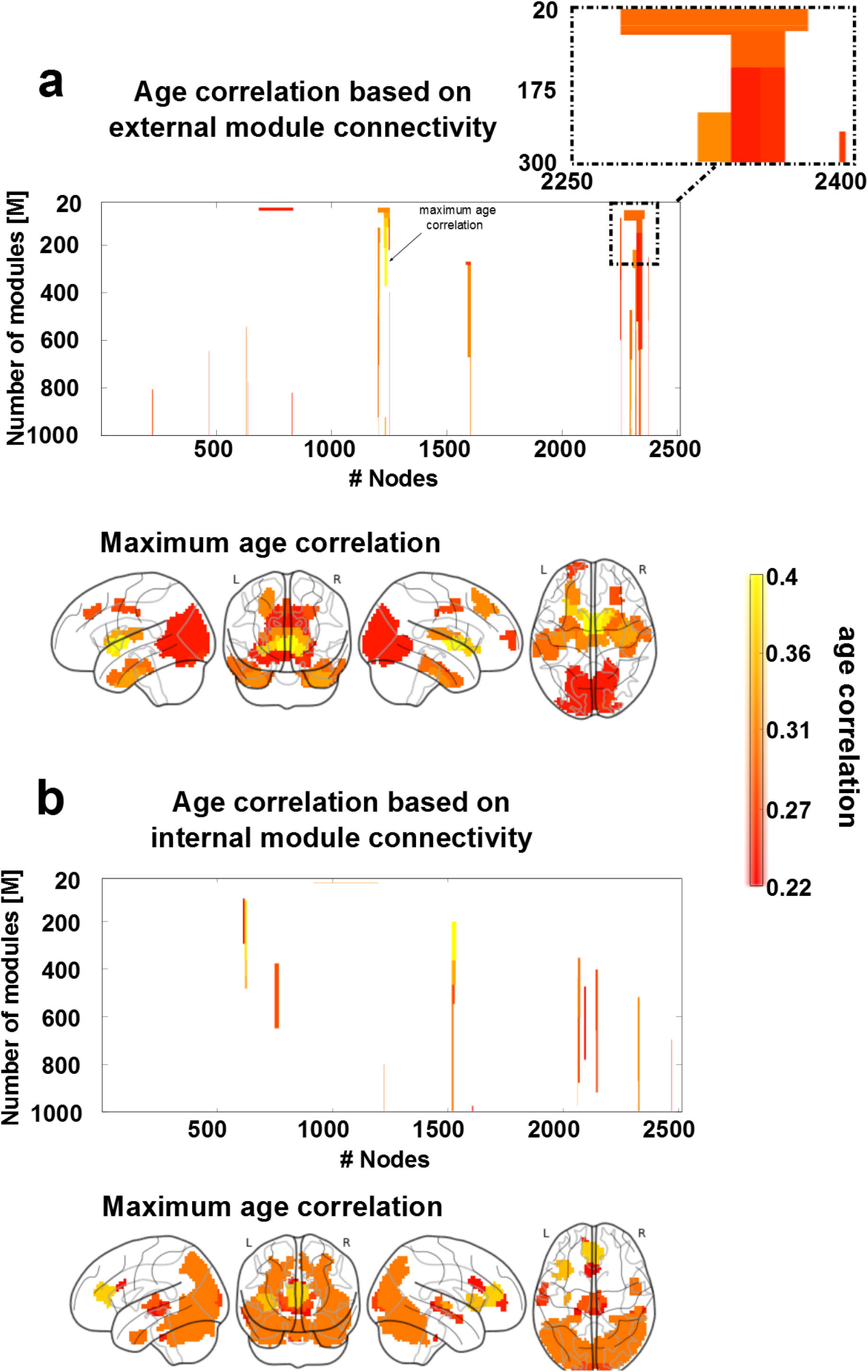
Structure-function correlo-dendrogram (CDG) and structure-function brain maps of age correlation across the multi-scale brain partition. To build structure-function age CDGs, we calculated for each module appearing in the BHA partition (20<=M<=1000) the correlation (and associated p-value) between age participant and FEC, FIC, SEC and SIC. **a:** Brain regions with external connectivity affected by age. 3D brains show per each of the 2514 regions, the value of maximum correlation achieved by that region among all values in the CDG (illustrated here with an arrow). A zoomed inset of the CDG is reported on the top right, showing how module division affects the correlation with age. **b:** Similar to panel a, but for age correlation with respect to internal connectivity.

*With regard to external connectivity* (figure 3a), maximum age correlations were found bilaterally in several cortical and subcortical AAL regions: the frontal superior and middle, cingulum middle, parahippocampus, calcarine, cuneus, lingual, occipital superior, middle and inferior, fusiform, precuneus, caudate, putamen, thalamus, temporal pole middle and temporal inferior.

*With regard to internal connectivity* (figure 3b), maximum age correlations were found bilaterally in the insula, cingulum anterior, calcarine, cuneus, occipital superior, middle and inferior, fusiform, parietal superior, angular, precuneus, thalamus, temporal middle and inferior and cerebellum.

Since the structure-function CDG does not provide any information on the individual contribution that either the structural or the functional descriptor has on the correlation value, we separated all possible four cases of combined correlations (increased structural and functional; decreased structural and functional; increased structural and decreased functional; decreased structural and increased functional) and reported them separately in figure 4.

**Figure 4.**
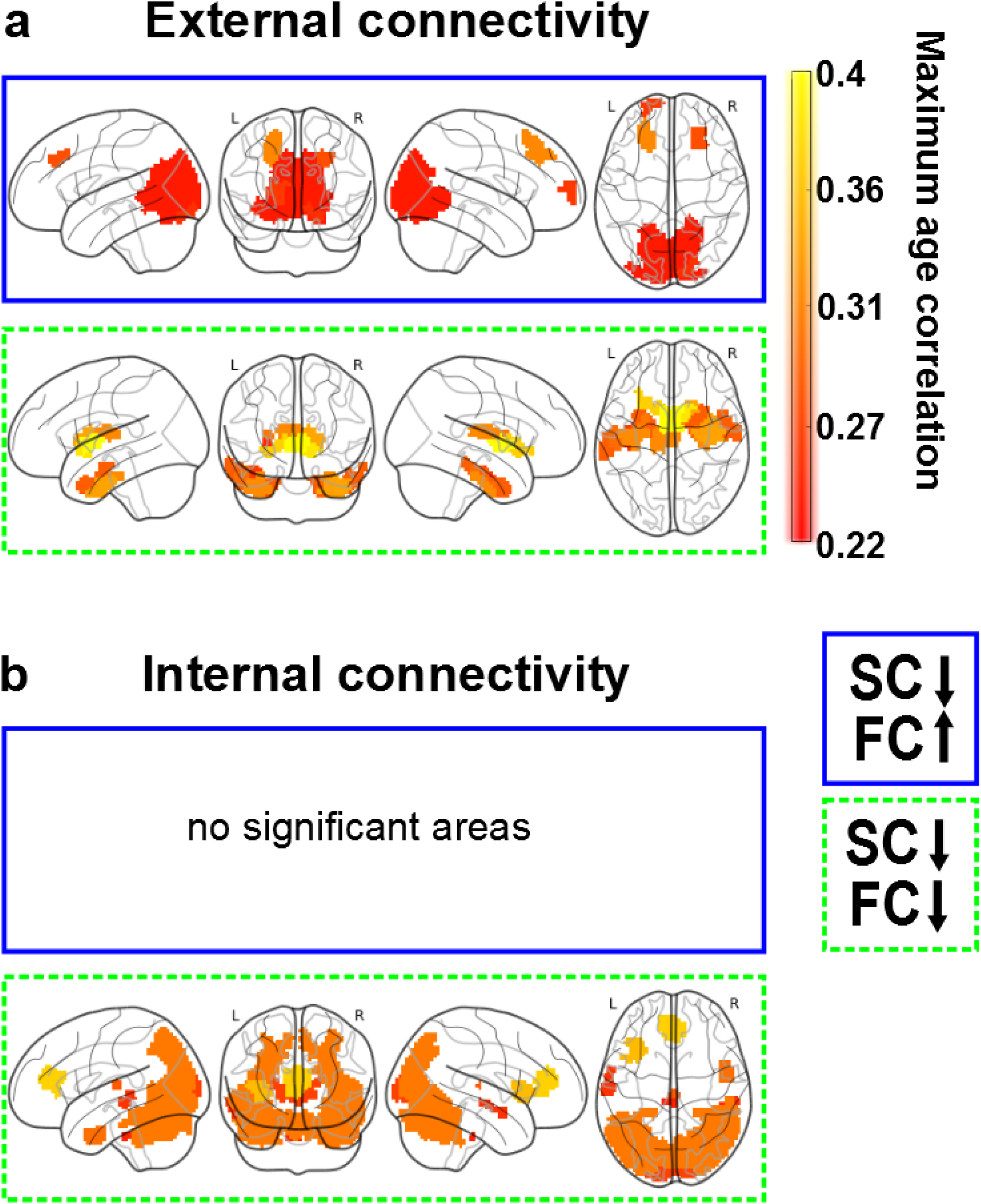
Functional connectivity modulation along lifespan. SC decreased with age, but FC might either increase (rectangle with solid blue line) or decrease (rectangle with dashed green line), and this occurred for both external connectivity descriptors (panel a) and internal connectivity descriptors (panel b). Notice that the situation of FC increasing and SC decreasing did not exist with regard to internal connectivity. As in figure 3, all the non-zero correlation values plotted here were statistically significant (after multiple-comparison correction).

The external connectivity analysis identified brain regions with *opposing* tendencies by which the structural connectivity decreased with age whilst the functional connectivity increased (figure 4a, blue rectangle), a mechanism representing *network homeostasis*. These regions were found bilaterally in the frontal superior and middle, calcarine, cuneus, lingual, occipital superior, middle and inferior and precuneus. Regions for which both structural and functional connectivities decreased with age (figure 4a, green rectangle) were found bilaterally in the parahippocampus, fusiform, caudate, putamen, thalamus and temporal pole middle and inferior.

The internal connectivity analysis did not identify any brain region in which the structural connectivity decreased with age and the functional one increased (figure 4b, blue rectangle), indicating that *network homeostasis* only was appreciated when looking to external connectivity patterns. Regions where both structural and functional connectivity decreased with age (figure 4b, green rectangle) were found in the insula, cingulum anterior (the anterior part of the default mode network), calcarine, cuneus, occipital superior, middle and inferior, fusiform, parietal superior, angular, precuneus, temporal middle and inferior and the cerebellum.

Next, we made use of the MLE to calculate BCA. We first pooled into a common list the four set of connectivity descriptors (FIC, FEC, SIC, SEC) and ordered them descendant in age correlation (that is, the first elements in the list were the descriptors that most correlated with age). By simply increasing the number of descriptors into the MLE, the correlation between ChA and BCA monotonously increased with age (figure 5a), that is, the more descriptors incorporated into the MLE the better the age estimation. This overfitting was resolved by splitting the entire dataset in two pieces, 75% of the dataset used for training (to calculate the MLE solution) and the remaining 25% used for testing (to calculate the MAE). By adding one by one descriptors each time and optimizing the regression performance (see Method), the *best* model in estimating age occurred for *K=38* descriptors, which provided a minimum mean MAE value (after U=100 repetition experiments) equal to 5.89 years (figure 5b). When calculating MLE using these K=38 best descriptors, the graphical representation of ChA as a function of BCA showed an excellent correspondence (figure 5c, correlation=0.95, p-value<10^-20^).

**Figure 5:**
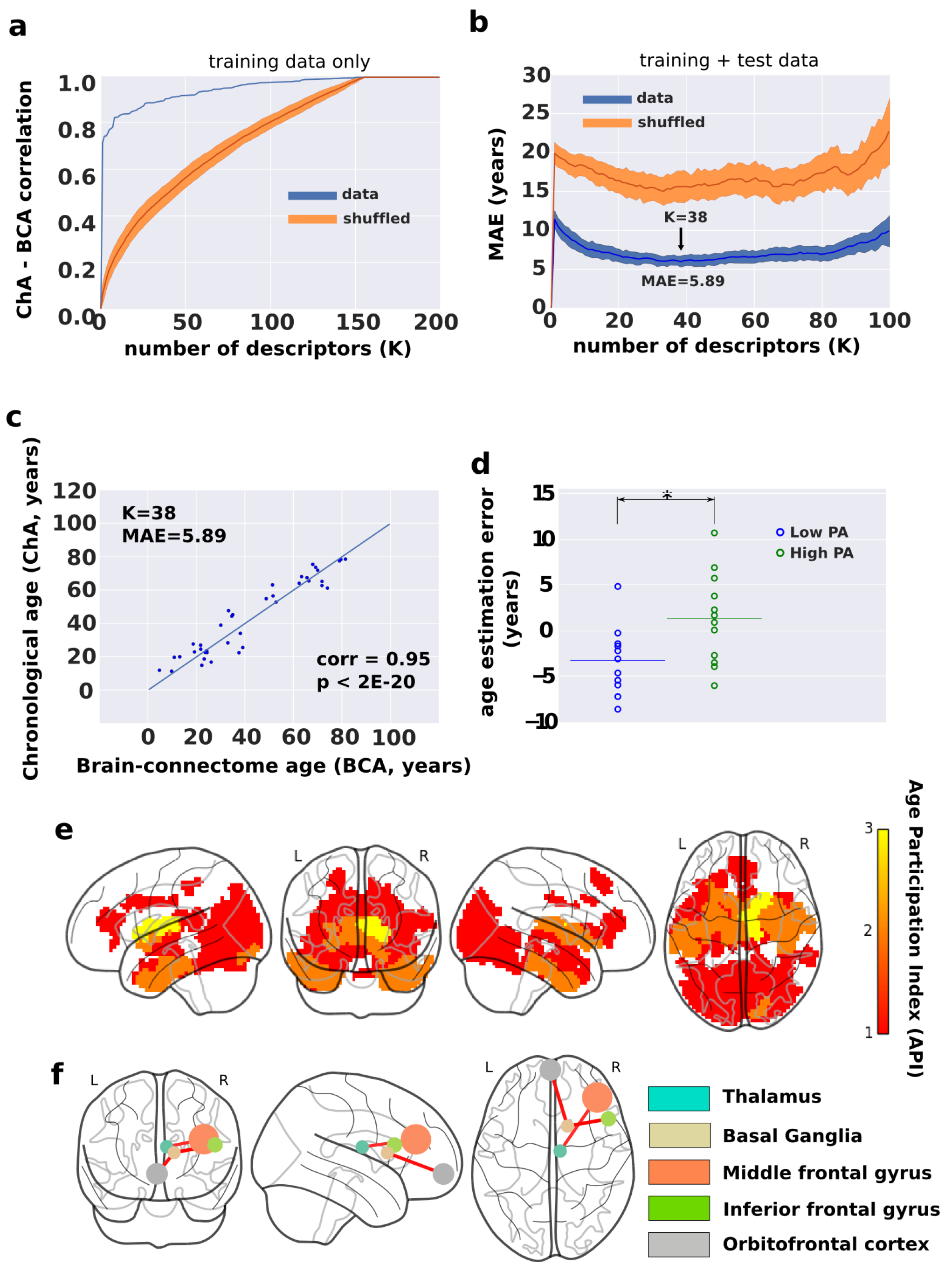
Chronological age (ChA) vs Brain-connectome age and its modulation by the amount of physical activity. **a:** Correlation between ChA and BCA as a function of the number of descriptors (K), when the entire dataset has been used for training (methods). Notice that, the larger number of descriptors, the higher the correlation. However, this strategy is well known to produce overfitting. In blue, we colored results from real data, and in orange, we plotted the results after shuffling the age vector a number of U=100 experiments, which provides the null-distribution (here represented the mean ± SD). **b:** Mean absolute error (MAE) as a function of K, using 75% of the dataset for training and 25% for testing. The minimum MAE, corresponding to 5.89 years provides optimal solution, achieved when K=38 different descriptors have been incorporated into the maximum likelihood estimator. Similar to panel a, blue and orange represent respectively real and shuffled data after Q=100 experiments. **c:** For a single estimate (chosen to have a similar MAE as the average one over the 100 experiments), we plot ChA (in years) as a function of the BCA (here, equal to the MLE solution with the best K=38 best connectivity descriptors), which provides a correlation value of 0.95 (p<2E-20). **d:** Age estimation error (defined as ChA minus BCA) for two groups of participants, one performing high physical activity (PA), and a different one with low PA values. **e:** Brain maps of the K=38 best descriptors. Color bar indicates age participation index (API), accounting for how many times one brain region is significantly correlated with age in relation to any of the four following categories: SEC, SIC, FEC and FIC. Basal ganglia and thalamus are the brain structures whose connectivity participates most prominently in ageing. **f:** Basal ganglia and thalamus connect according to a structure-function manner to the inferior and middle frontal gyri together with the orbitofrontal cortex, ie. the so called fronto-striato-thalamic (FST), is the major pathway participating in brain aging. Node size is proportional to the volume size of the region that participates in this network, whereas link thickness is proportional to structure-function correlation values (Table 1).

Next, we obtained brain maps of the best *K=38* descriptors (figure 5e). Because by construction of our model each of the 2514 regions might participate in four different classes of descriptors (FIC, FEC, SIC, SEC), we plotted each region according its *Age Participation Index (API),* an integer number between 0 and 4 and indicating with how many of the four classes a specific brain region contributed to the age estimator model.

Regions with API equal to 1 were found bilaterally in the frontal superior, insula, cingulum anterior and middle, hippocampus, parahippocampus, calcarine, cuneus, lingual, occipital superior, middle and inferior, fusiform, precuneus, thalamus, temporal middle and inferior and temporal pole middle. Regions with API equal to 2 were found bilaterally in the parahippocampus, fusiform, caudate, putamen, thalamus, temporal pole middle and temporal inferior. Finally, regions with API equal to 3, and therefore, the regions with a major correlate of physiological ageing were found in the connectivity of basal ganglia (caudate, putamen, pallidum) and thalamus.

Next, we looked into the networks that starting from regions with API=3 were connected to the rest of the brain. These regions were connected functionally and bilaterally to orbitofrontal (superior, middle and inferior), middle frontal, olfactory, gyrus rectus, cingulum (anterior, middle and posterior), calcarine, occipital middle, fusiform, precuneus, temporal middle and cerebellum. Structurally, these regions were inter-connected and also with the insula. Finally, regions with API=3 (including part of the caudate, putamen, pallidum and thalamus) were structurally-functionally connected to the orbitofrontal cortex and to the inferior and middle frontal gyri (figure 5f). The volume values within each brain structure participating in this circuit together with the connectivity values (link strength) between these structures are given in Table 1.

**Table 1.** Structure-function connectivity from basal ganglia and thalamus to orbitofrontal and frontal areas is the key circuit for brain ageing. Volume sizes within each region participating in this network together with link strength (ie. structure-function significant correlation value) between pair regions. X represents no existing links.

|  | Thalamus<br>(2970 mm<sup>3</sup>) | Middle Frontal<br>Gyrus<br>(16443 mm<sup>3</sup>) | Basal Ganglia<br>(2727 mm<sup>3</sup>) | Inferior Frontal<br>Gyrus<br>(3942 mm<sup>3</sup>) | Orbitofrontal<br>Cortex<br>(8100 mm<sup>3</sup>) |
| --- | --- | --- | --- | --- | --- |
| Thalamus<br>(2970 mm<sup>3</sup>) | x | 0.238 | x | x | x |
| Middle Frontal<br>Gyrus<br>(16443 mm<sup>3</sup>) | 0.238 | X | x | x | x |
| Basal Ganglia | x | X | x | 0.285 | 0.27 |
| (2727 mm<sup>3</sup>) |  |  |  |  |  |
| Inferior Frontal Gyrus<br>(3942 mm<sup>3</sup>) | x | X | 0.285 | x | x |
| Orbitofrontal Cortex<br>(8100 mm<sup>3</sup>) | x | X | 0.27 | x | x |

We finally asked whether physical activity (PA) can in any manner modulate BCA (ie., if PA can modify the brain’s structural-functional wiring in a manner that age estimation has a systematic bias). To provide an answer to that question, we compared the estimation error (defined as ChA minus BCA) in two groups, one group with high values of PA (90^th^ percentile) and another one with low values of PA (10^th^ percentile, figure 5d). By building the MLE with the best *K=38* descriptors, we obtained a mean estimation error of -3.2 years (SD=-3.7) for the low PA group, and of 1.33 years (SD=4.94) for the group with high PA. The estimation error was significantly different in the two groups (p-value=0.02 after a Kolmogorov–Smirnov test), and this occurred independently of ChA, as the correlation value between the estimation error and age was -0.05 (p-value=0.88) for the group with high PA and 0.39 (p-value=0.23) for the group of low PA.

## Discussion

The chronological age (ChA) differs from the biological one. Whilst the former is defined as the time running since birth, the latter quantifies the maturity level that an individual (or an organ) has at the operational level. In relation with the brain, the discrepancy between the brain age and ChA might constitute a biomarker for quantifying deterioration as a result of disease or improvement after some treatment or therapy, which has unlimited applications. Here, we asked whether the brain age could be determined exclusively based on structure-function connectivity descriptors, coined as brain-connectome age (BCA). Therefore, we did not take into consideration typical morphological descriptors (such as grey and white matter atrophy or ventricular volume) that have been widely shown to correlate with ChA. Our results demonstrate that BCA can estimate ChA with high accuracy, determined by a mean absolute error of 5.89 years in a group of N=155 participants with age ranging from 10 to 80 years. Our results also reveal that the basal ganglia/thalamus and their connection with orbitofrontal and frontal areas is the key circuit accounting for brain ageing. Furthermore, we also report that the level of physical activity (PA) rejuvenates the brain connectome, implying that high-PA participants have an associated BCA that qualifies them as being younger than low-PA participants.

## Differences between brain age and chronological age by assessing brain morphology

Several studies have assessed discrepancies between brain morphology age and ChA as an estimation of brain functioning in pathological groups. In relation with mild cognitive impairment (MCI), it was shown that the brain age could become even 10 years higher than ChA^56^. In a different study, it was shown that the error in age estimation predicted the conversion from MCI to Alzheimer’s disease better than any other variable ^40^, as compared to imaging morphological variables (such as the volume of subcortical structures), cognitive scales or protein biomarkers in cerebrospinal fluid. One-year bias was associated to 10% higher risk conversion. In relation to other pathologies, and also using morphological descriptors to estimate the brain age, the differences between brain age and ChA explained for instance brain deterioration from attenuated psychosis to chronic schizophrenia^54^, brain deterioration in patients with human immunodeficiency virus^41^, accelerated atrophy after traumatic brain injury^39^ (suggesting that the chronic effects after the insult can resemble normal ageing), but also brain rejuvenation after meditation^58^.

Morphological age-related alterations have been reported in all body organs, such as liver, kidney, heart, lung, skin, but notably, what makes the brain distinct from other organs is precisely its complex wiring, where short-range connectivity operates at multiple scales in combination with long range circuitry, allowing the two main brain functional principles of segregation and integration^77^ to work in harmony. Therefore, the BCA model presented in this work provide a new complementary and fundamental approach within the above framework, being fully centred on the multi-scale organization of the brain circuitries/networks, therefore correlating ageing with lower and higher brain functions.

## Differences between brain age and chronological age by assessing brain connectivity: The importance of combining SC and FC

Only very few studies have made use of connectivity metrics for age estimation, but none of them combined SC and FC metrics to perform the estimation. By combining morphological descriptors together with functional connectivity ones, Liem et al (2017) were able to improve age estimation up to 4.29 years^53^. In relation to connectivity metrics, it was shown in a seminal study that resting FC descriptors estimated brain age^51^, but rather than addressing physiological ageing, the authors focused on neural development in the age range between 7 and 30 years. By using only structural networks, but not functional data, it was shown that a simple metrics such as the sum of all connectivity links (ie., streamline number) weighted by the age-link correlation, estimated ChA with high precision^52^ in healthy participants aged 4 to 85 years.

By combining both SC and FC descriptors in the present study, we have achieved a high performance, quantified by a MAE equal to 5.89 years. To the best of our knowledge, such an approach has never been reported before in the context of age-estimation. When repeating the entire procedure using only SC descriptors, the performance was worse (MAE=8.1 years). Analogously, when using only FC descriptors, the performance was also worse (MAE=8.45 years).

Our results not only point out the relation between BCA and the aging process, but also highlight how the wiring and the dynamics of the brain networks cannot be disentangled without losing the emergent synergetic picture of their operational complexity. Indeed, when looking solely at function, as it happens in task fMRI, a distinct scenario emerges, the so-called *frontal super-activation*, where younger people exhibit no or lateralized frontal activation when performing the task, while older people incorporate unilateral or bilateral frontal cortex activation^78^. Our BCA model, although obtained when the brain is at rest, extends task-related studies, revealing that the fronto-subcortical (striatum and thalamus) complex is the primary circuit critically accounting for the functional age-deterioration. Indeed, the key element is that the multi-scale modular BCA approach used in this work perfectly preserves the intrinsic link between structure and function. In fact, the resting activity is shaped by modular organization, where the structural and functional architecture clearly merge and match each other^64^. Therefore the multi-scale modular BCA approach used in this work perfectly preserves the intrinsic link between structure and function.

## The structure-function connectivity between subcortical areas (striatum and thalamus) and frontal/orbitofrontal cortex, mediating cognitive control functions, is the principal determinant of brain ageing

Several studies have shown that the connectivity profile of the basal ganglia (BG) and thalamus is affected by ageing and correlates with age-related neuropsychological decay. For example, a reduction in thalamic volume along the lifespan has been associated with age-related sensorimotor performance deterioration^79^. Beyond the classical relation to motor function^80^, there is nowadays mounting evidence to associate the BG decline with executive function deficits along the lifespan, such as motor switching^81^ and inhibitory control^82,83^, but also in motor learning^84^, whereby older adults perform worse than young. The status of connectivity between the thalamus and BG, by means of the fronto-striato-thalamic (FST) circuit, has been associated with task-switching performance^85–88^ which measures the suppression of certain actions to flexibly adapt to new different ones. These so-called *cognitive control* functions are altered along the lifespan^7^. On top of these results, and without a priori assuming the participation of these regions, we have provided quantitative evidence that the FST circuit delivers the major contribution for age estimation.

## Effects of physical activity on the brain connectome age

As far as we know, our study is the first one demonstrating that physical activity (PA) can rewire the human connectome to increase its resilience by slowing down brain ageing. In relation to brain morphology, it has been shown^57^ that none of the following activities have a strong implication in the estimation of brain age: walking/hiking, jogging, running, bicycling, aerobic exercise, lap swimming, tennis/squash/racquetball or low intensity exercise, but in addition, the same authors showed a small association between brain age and a variable named “*daily number of flights of stairs climbed”*. Here, by only looking to structure-function connectivity descriptors rather than morphological variations, we showed strong evidence between PA and ChA estimation. Notably, and distinct from previous work^57^, our methodology does not need to disentangle which specific type of physical exercise significantly impacts brain aging, although it has the potentiality.

## A novel methodology to quantify the structure-function brain connectivity impact of therapies, diseases and lifestyles

The novelty of our approach is based on identifying, on a multi-scale level, brain modules whose connectivity correlate structurally and functionally with age. The significant descriptors are then pooled to create an optimized linear regression model capable to assess the biological brain age of a given connectome with minimal error. Such a methodology opens two important and different perspectives. First, it provides a quantitative approach to assess the impact of therapies on biological brain ageing (ideally rejuvenating the brain connectome), diseases (accelerating the connectome ageing) and other factors shaping lifestyle. Secondly, the same methodology can be used to correlate any graded variable (not only age) to the brain connectome, allowing for brain-connectome estimators of any other biomarker. In this framework, a biomarker would be any functional, structural or behavioural score measured in participants, and the brain-connectome linear model developed in this work will be first used to fit participant scores and next, to estimate new single subject scores. Lastly, our method makes use of network node metrics to estimate brain age, but this method can also be extended to network link level, thus identifying specific pathways rather than brain regions, provided a large cohort is available to control for multiple comparisons (as a given connectome has N nodes and N^2^ links).

## Summary

We have shown that the chronological age can be estimated from the structural-functional connectome with a much higher accuracy than structural or functional connectivity separately. Using the mismatch between chronological and brain age might be useful for quantifying the brain’s deterioration or reorganization after new treatments, implying a multitude of meaningful applications. In particular, we have shown that participants who exhibit higher levels of physical activity have a brain connectome age that appears younger than the chronological one, but also that low levels of physical activity increase the brain connectome age. Therefore, our results demonstrate that physical activity makes the brain more resilient by slowing brain ageing. By using a blind approach in which no brain structures were a priori assumed to be affected by ageing, our method has shown that the connectivity of the fronto/orbitofrontal-striato-thalamic circuit is critically important for brain ageing, consistent with previous work associating this circuit to age-related deterioration of cognitive control of action.

## Author Contributions

PB and AE had equal first-author contribution; MBP, LP and SPS collected the MRI data and physical activity scores; ID preprocessed the MRI data; PB, AE performed the analysis; PB, AE and JMC made the figures. All the authors wrote the manuscript and agreed with its submission; SPS and JMC had equal last-author contribution.

## Acknowledgements

JMC and PB acknowledge financial support from Ikerbasque (The Basque Foundation for Science) and from the Ministerio Economia, Industria y Competitividad (Spain) and FEDER (grant DPI2016-79874-R to JMC, grant SAF2015-69484-R to PB). AE acknowledges financial support from the Basque Government (Eusko Jaurlaritza), grant PRE/2014/1/252. IG acknowledges financial support from Instituto de Salud Carlos III “JR15/00008” (co-funded by European Regional Development Fund/European Social Fund, “Investing in your future”). MPB is supported by post-doctoral fellowship and a research grant of the Research Foundation Flanders (FWO, grant 1504015N). SPS is supported by KU Leuven Special Research Fund (grant C16/15/070) and the Research Foundation Flanders (FWO, grant G0708.14).

